# The prevalence and benefits of admixture during species invasions: a role for epistasis?

**DOI:** 10.1101/139709

**Authors:** Brittany S Barker, Janelle E Cocio, Samantha R Anderson, Joseph E Braasch, F Alice Cang, Heather D Gillette, Katrina M Dlugosch

**Affiliations:** Department of Ecology & Evolutionary Biology, University of Arizona, Tucson, AZ, USA

**Keywords:** multiple introductions,, invasiveness,, heterosis,, heterozygosity-fitness correlations,, genetic diversity,, epistasis

## Abstract

Species introductions often bring together genetically divergent source populations, resulting in genetic admixture. This geographic reshuffling of diversity has the potential to generate favorable new genetic combinations, facilitating the establishment and invasive spread of introduced populations. Observational support for the superior performance of admixed introductions has been mixed, however, and the broad importance of admixture to invasion questioned. Under most underlying mechanisms, admixture’s benefits should be expected to increase with greater divergence among and lower genetic diversity within source populations. We use a literature survey to quantify the prevalence of admixture and evaluate whether it occurrs under circumstances predicted to be most beneficial to introduced species. We find that 39% of species are reported to be admixed when introduced. Admixed introductions come from sources with a wide range of genetic variation, but are disproportionately absent where there is high genetic divergence among native populations. We discuss multiple potential explanations for these patterns, but note that negative epistatic interactions should be expected at high divergence among populations (outbreeding depression). As a case study, we experimentally cross source populations differing in divergence in the invasive plant *Centaurea solstitialis*. We find many positive (heterotic) interactions, but fitness benefits decline and are ultimately negative at high source divergence, with patterns suggesting cyto-nuclear epistasis. We conclude that admixture is common in species introductions and often happens under conditions expected to be beneficial to invaders, but that these conditions may be constrained by predictable negative genetic interactions, potentially explaining conflicting evidence for admixture's benefits to invasion.

## Introduction

Introduced non-native species are now a common feature of ecosystems on Earth, and they are recognized as being one of the dominant sources of biodiversity change in the Anthropocene (Ellis *et al.* 2012; Vellend *et al.* 2013). While many factors will shape the success of species introductions and their ecological interactions with recipient environments (Sakai *et al.* 2001), there is increasing evidence that genetic factors can play a role in this process (Baker & Stebbins 1965; Ellstrand & Schierenbeck 2000; Lee 2002; Cox 2004; Colautti & Barrett 2013; Rius & Darling 2014; Whitney & Gering 2015; Mesgaran *et al.* 2016). An understanding of when, where, and how genetic changes influence the outcomes of colonization should inform broader questions about when species introductions will lead to establishment and invasive spread (Hufbauer 2008, 2017; Lee & Gelembiuk 2008; Forsman 2014; Rius & Darling 2014; Szűcs *et al.* 2014, 2017; Bock *et al.* 2015; Colautti & Lau 2015; Dlugosch *et al.* 2015a; Williams *et al.* 2016; Ochocki & Miller 2017; Weiss-Lehman *et al.* 2017).

In particular, successful introductions are often the products of multiple inputs of genetic material from different source populations, resulting in the admixture of divergent genotypes (Dlugosch & Parker 2008a; Uller & Leimu 2011; Dlugosch *et al.* 2015a). Admixture has the potential to facilitate species invasions by creating unique opportunities for positive genetic interactions among previously isolated alleles and for adaptive evolution of novel genotypes, which could increase the fitness of admixed populations (Kolbe *et al.* 2004; Lavergne & Molofsky 2007; Keller & Taylor 2010; Wagner *et al.* 2017). These same mechanisms are known to have contributed directly to non-native species establishment and the rise of particularly invasive novel genotypes in cases involving hybridization between species, and the potential for similar benefits of admixture within species appears widespread (Ellstrand & Schierenbeck 2000; Drake 2006; Hovick & Whitney 2014). Consequently, there is intensifying interest in the potential for genetic admixture to provide a general mechanism by which many non-native species are able to establish and develop into invaders (Frankham 2005; Hufbauer 2008, 2017; Verhoeven *et al.* 2011; Molofsky *et al.* 2014; Rius & Darling 2014; Bock *et al.* 2015; Dlugosch *et al.* 2015a).

Empirical support for the role of admixture in facilitating invasions has been mixed, however. On one hand, positive correlations between evidence of admixture and fitness traits have been identified in some invasions (Keller *et al.* 2014; Rius & Darling 2014), and experimental admixtures have performed better in several studies (Turgeon *et al.* 2011; van Kleunen *et al.* 2015; Wagner *et al.* 2017). On the other hand, studies that have found no association between admixture and increased invasiveness have called into question whether mixing of divergent populations can realistically be expected to contribute to increased fitness and introduction success across many invaders (Wolfe *et al.* 2007; Dutech *et al.* 2012; Chapple *et al.* 2013). These conflicting results are not necessarily surprising given that studies of native species have long demonstrated that mating across different populations can have fitness effects ranging from positive (i.e.alleviating inbreeding depression) to detrimental (i.e. resulting in outbreeding depression), depending upon the mechanisms underlying the genetic interactions (Price & Waser 1979; Lynch 1991; Edmands 1999; Keller & Waller 2002; Reed & Frankham 2003; Birchler *et al.* 2010; Frankham *et al.* 2011; Chen 2013)

Given that introduced species are often derived from multiple source populations, genetic admixture could be a frequent path to the evolution of invasiveness, but only if its fitness effects are typically positive under conditions commonly experienced during introductions. There are several non-mutually-exclusive mechanisms that could generate positive fitness effects as a consequence of either the increase in genetic diversity that should result from combining divergent populations, or the genetic interactions that can result from bringing together new combinations of alleles within individuals (Lynch 1991; Dlugosch *et al.* 2015a; Hufbauer 2017):

- *Evolutionary Rescue*. Populations that will go extinct or fail to spread because they lack adaptation to local conditions could be rescued by inputs of additional genetic variation, a scenario known as ‘evolutionary rescue’ (Carlson *et al.* 2014). For introduced species in particular, additional inputs of genetic diversity could facilitate both adaptation to the novel environment of introduction and adaptation to colonization in general, such as with increases in dispersal ability and expansion rate (Thompson 1998; Cox 2004; Holt *et al.* 2005; Phillips *et al.* 2006; Prentis *et al.* 2008; Weiss-Lehman *et al.* 2017). Evolutionary rescue should be most likely in populations with low genetic diversity (such that adaptive variation is limiting), and increasingly impactful as the divergence between admixing populations increases, allowing the combination of greater numbers of alleles (Rieseberg *et al.* 2007; Wagner *et al.* 2017; Ochocki & Miller 2017; Szűcs *et al.* 2017).
- *Complementarity*. As diversity increases within a population, different genotypes may occupy somewhat different and complementary niches, sometimes increasing the mean fitness across the population as a whole (Crawford & Whitney 2010; Chen *et al.* 2015). Complementarity should be most beneficial at low genetic diversity within a focal population, where niche diversity among genotypes is low, and become increasingly likely as divergence between admixing populations increases and greater numbers of genotypes are combined (Wang *et al.* 2012; Le Roux *et al.* 2014), though this effect may plateau after niches are exhausted (Ellers *et al.* 2011).
- *Genetic Rescue*. Also known as ‘directional dominance’ (Birchler *et al.* 2010; Chen 2013), genetic rescue refers to the rescue of deleterious inbred (homozygous) loci by outbreeding with a divergent population (Tallmon *et al.* 2004). Homozygous loci in introduced populations can be derived from both historic genetic load already present in native source population, and additional fixation of deleterious variants during founding events (Excoffier *et al.* 2009). Multiple introductions from divergent sources can therefore provide genetic rescue to an establishing population by contributing/restoring superior alleles that increase the fitness of introduced genotypes, potentially resulting in superior genotypes that transgress the fitness of both parental populations (Ellstrand & Schierenbeck 2000; Hufbauer 2008; Keller & Taylor 2010; Rius & Darling 2014; van Kleunen *et al.* 2015). Genetic rescue should benefit invasions when diversity within founding populations is low and there is significant genetic load to rescue (Lynch *et al.* 1995; Lohr & Haag 2015). The fitness benefits should scale positively with divergence between source populations, until all loci are rescued and fitness gains plateau (Lynch1991).
- *Overdominance*. Also known as ‘heterozygote advantage’, overdominance occurs when heterozygous allele combinations have higher fitness than any homozygous genotype (Birchler *et al.* 2010; Chen 2013). These effects are expected to manifest primarily in the first (F1) generation of an admixture event, but decay quickly due to the increase in homozygosity that occurs over subsequent generations, unless heterozygosity can be preserved by asexual propagation, polyploidy, or other means (Ellstrand & Schierenbeck 2000; Drake 2006; Facon *et al.* 2008). These effects should be strongest when opportunities for novel heterozygosity in admixed progeny are highest, i.e. when there are greater numbers of fixed differences between source populations.
- *Epistasis*. Epistasis occurs when alleles at different loci interact (Phillips 2008). Epistatic interactions that arise from admixture are predicted to have increasing fitness effects as divergence among source populations increases, due to natural selection for locally co-adapted allele combinations and/or to the build up of Bateson-Dobzhansky-Muller incompatibilities from genetic drift, acting separately in each source population (Lynch & Walsh 1998; Moyle & Nakazato 2010). The fitness effects of epistatic interactions in the first generation can be positive (heterotic) but will become increasingly negative with greater genetic distance between parents, particularly in later generations when co-evolved multi-locus genotypes are broken apart by recombination (i.e. ‘hybrid breakdown’; Lynch 1991; Orr & Turelli 2001; Bomblies *et al.* 2007). Negative epistatic effects are thought to be one of the most important paths to reproductive isolation and speciation, and could impede admixture of divergent sources during multiple introductions, though the influence of these effects on species invasions is rarely discussed and largely unknown (Carroll *et al.* 2003; Dlugosch *et al.* 2015a).

Based on these mechanisms, fitness benefits from admixture should be expected to vary in predictable ways (Fig. 1a,b). As divergence between sources increases, opportunities for evolutionary rescue, complementarity, rescue of genetic load, the creation of overdominant heterozygotes, and epistatic interactions among loci should all increase. These interactions should all be positive for fitness initially, though benefits of most effects should ultimately plateau, and epistatic interactions will become increasingly negative with increasing divergence (Fig. 1a). Benefits of all mechanisms other than epistasis should also be strongest where founding populations harbor low within-population diversity (Fig. 1b), especially to the extent that this represents fixation of deleterious alleles (Lohr & Haag 2015) [with the caveat that species with a history of inbreeding may have purged genetic load (Crnokrak & Barrett 2002) and therefore stand to benefit only from evolutionary rescue, complementarity, and/or overdominance when at low genetic variation]. Thus the fitness effects of admixture should be expected to vary in magnitude under different scenarios, but to be either positive or neutral under most mechanisms (other than epistasis), consistent with the idea that admixture could be almost universally beneficial to invaders.

**Fig 1.**
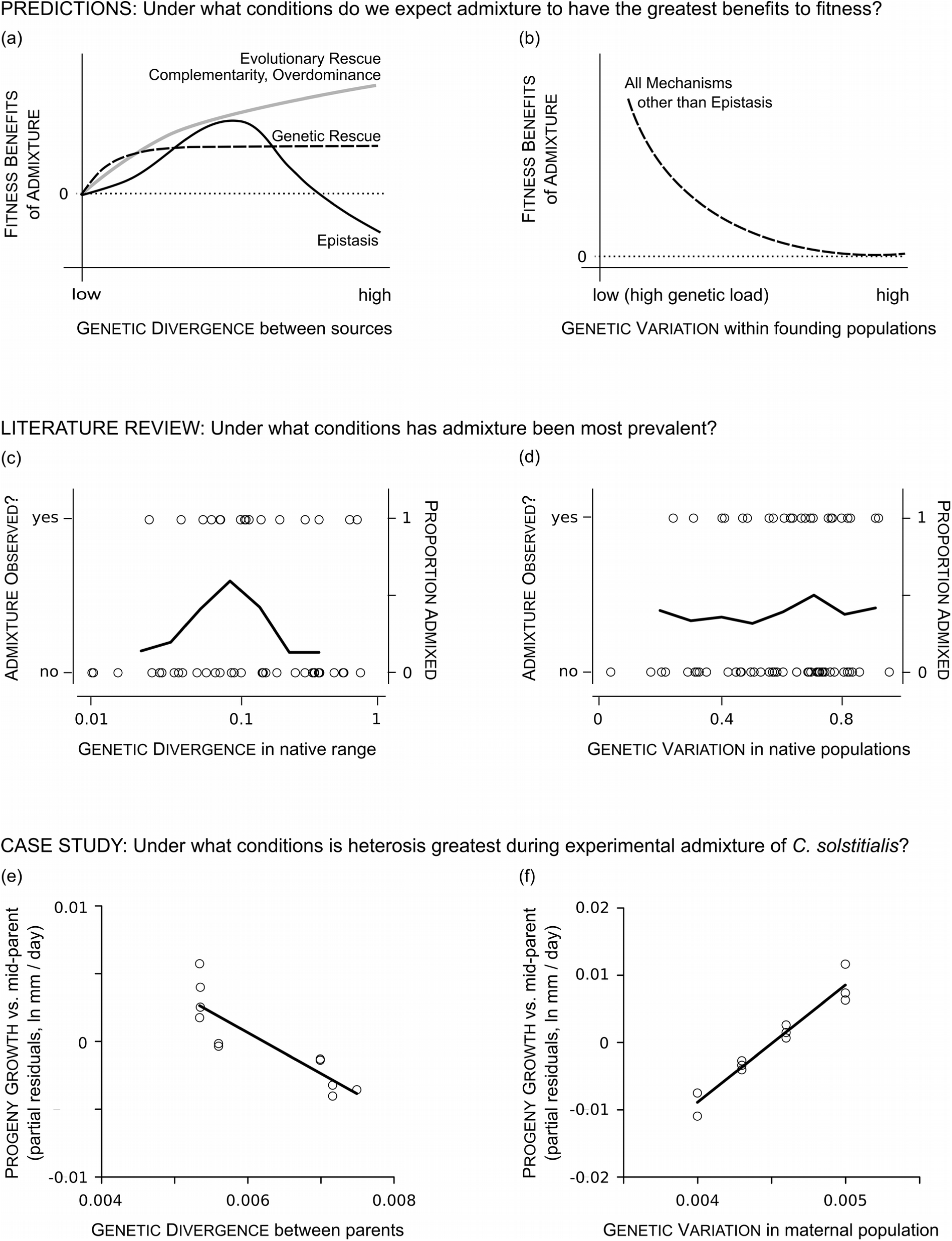
Identifying the conditions under which admixture might be most favorable during invasions. Shown are predicted relationships for the performance of admixed populations as a function of (a) the genetic divergence between their source populations and (b) the genetic variation within their initial founding populations, under different genetic and evolutionary mechanisms. In the literature, genetic admixture is reported more often at intermediate genetic divergence (FST and related metrics for microsatellite markers) among potential source populations in the native range (c), and shows no relationship with average genetic variation (HE) within native populations (d). In a linear model of progeny performance in experimental crosses the invasive plant *C. solstitialis* (excluding heterotic outlier [Asia × southern Greece]), genetic divergence (interpopulation π) had a significant negative effect (e, *P* < 0.01) and genetic variation (intrapopulation π) in maternal populations had a significant positive effect (f, *P* < 0.001) on the deviation of progeny from mid-parent genotype expectations. Data points in (e,f) show progeny deviation from mid-parent as partial residuals after taking into account all other effects in the model.

So where do known admixture events in introduced species fall in the parameter space of divergence among source populations and genetic variation within founding populations? To date there has been little study of the divergence among potentially admixing populations during species introductions (Dlugosch *et al.* 2015a). The question of how much genetic diversity is available in introduced populations has received considerably more attention. Previous studies have shown that introduced populations generally do not experience large reductions in diversity relative to native populations (though certainly many exceptions exist) and that low levels of diversity do not prevent adaptation (Dlugosch & Parker 2008a; b; Uller & Leimu 2011; Szűcs *et al.* 2017). Nevertheless, a lack of strong founder/bottleneck effects do not preclude the presence of historical genetic load and/or low diversity derived from source populations. Manipulations of genetic diversity have shown a range of effects on the performance of experimental invading populations, and the potential benefits of admixture in this regard remain an active area of research (Crawford & Whitney 2010; Szűcs *et al.* 2014, 2017; Williams *et al.* 2016; Hufbauer 2017; Wagner *et al.* 2017; Ochocki & Miller 2017; Weiss-Lehman *et al.* 2017).

In this study, we survey the literature to determine how native populations of successfully introduced species vary in their interpopulation divergence and intrapopulation diversity, and how these genetic features relate to observed instances of admixture in introductions. We ask whether admixture is being realized under conditions in which we might expect it to be favorable to many invaders. As a case study, we experimentally test for the fitness effects of genetic interactions in controlled crosses of *Centaurea solstitialis* L. (‘yellow starthistle’; Asteraceae), a highly invasive annual plant in the Americas. We cross native populations that span a range of genetic divergence and diversity to test for associations between the fitness of admixed progeny and these factors. Using the results from both our literature and experimental analyses, we consider the likelihood that admixture is a general mechanism promoting the invasiveness of introduced species.

## Materials and Methods

### Literature Review

Using papers that reported the distribution of molecular genetic variation in native populations and in populations introduced from the native range to new areas (i.e. ‘primary’ introductions), we extracted metrics of genetic variation and divergence for native populations and recorded whether admixture was reported in the introduced populations. To identify these papers, we revisited all studies in several reviews of genetic variation in introductions (Roman & Darling 2007; Dlugosch & Parker 2008a; Uller & Leimu 2011), as well as all papers citing these reviews on Web of Science and Google Scholar through June 2014. We used the most recent or comprehensive analysis for a given species when multiple analyses were available.

We relied on the judgement of the authors for determinations of admixture, but noted the primary basis for the determination. These included 1) formal admixture tests among alternative models of historical demography (e.g. using Approximate Bayesian Computation); 2) formal population genetic admixture tests (e.g. based upon expected deficiencies of heterozygosity or elevated LD); 3) patterns of elevated genetic diversity in introduced populations relative to all sampled native populations; 4) similarity of invader genotypes to potential native source populations (e.g. phylogenetic trees, haplotype networks, distance trees, ordination, and assignment tests); and 5) population structure analyses, where admixture is inferred from the inclusion of individuals in more than one inferred population (an approach that has recently been shown to be prone to incorrect assignment of admixture under many scenarios; Falush *et al.* 2016; Novembre 2016; Wang 2017) Where determination of admixture was ambiguous for the authors, we removed the study from our statistical analyses. Where introduction events of the same species into different regions included admixture in some areas and no admixture in other areas, the species was coded as admixed for analysis.

Using a similar survey, we previously reported a relationship between genetic divergence among potential native source populations and reporting of admixture in their introductions (Dlugosch *et al.* 2015a; Dryad doi:10.5061/dryad.s2948). This study expands on those data and presents several new analyses. We first tested for the possibility that sampling design affected reports of admixture and its relationship with genetic divergence or diversity, including effects of sample size, the number of sites sampled, and marker type. For the most commonly used marker in our survey (microsatellites, see Results), we quantified potential source population divergence using FST (and related metrics) reported among all native populations in a study, and tested for a relationship with admixture using logistic regression. We also tested for the effect of genetic variation within native populations on the likelihood of reporting admixture in an introduction using logistic regression. We quantified genetic variation as the mean across all native sampling sites and loci (within a single type of genetic marker) of expected heterozygosity (HE) and observed heterozygosity (HO). All statistical tests were performed in JMP 11 (SAS Institute, Cary, USA).

### Case Study: Centaurea solstitialis

We experimentally tested for genetic interactions in controlled crosses among native populations of *C. solstitialis* spanning a range of genetic diversity and divergence. This species was introduced in large numbers to the Americas as a seed contaminant of alfalfa stock imported from the Old World, where it escaped agricultural fields and became a major pest of grasslands (Gerlach 1997). A lineage in western Europe appears to be the product of ancient admixture between populations from eastern Europe and Asia, and has served as a bridgehead for invasions in the Americas (Barker *et al.* 2017). Several invading populations show evidence of additional recent admixture with other native populations (Dlugosch *et al.* 2013; Eriksen *et al.* 2014; Barker *et al.* 2017). Phenotypic studies have revealed evolutionary increases in plant size during this range expansion, which is associated with higher fitness (Widmer *et al.* 2007; Eriksen *et al.* 2012; Dlugosch *et al.* 2015b). Experimental crosses in *C. solstitialis* provide an opportunity to better understand how admixture might be influencing the evolution of invasiveness.

### Collections

Genotypes of *C. solstitialis* were collected from 21 native sites (Supporting Information Table S1) spanning multiple potential source regions across western and eastern Europe, Asia, and southern Greece (Gerlach 1997; Tutin *et al.* 2010; Barker *et al.* 2017). Seeds were collected from wild plants during August–September 2008, from each of 9−22 maternal plants located at least1 m apart along a linear transect at each site. In Asia, seeds of mothers at each site were combined into bulk collections by site. The species was identified by the authors according to the Flora Europaea (Tutin *et al.* 2010) and vouchers from each sampling site are available at the University of Arizona herbarium (ARIZ, accession numbers in Supporting Information Table S1).

### Genetic variation within and divergence among native populations

We previously identified four geographically-structured, genetically-divergent populations in the native range (see Fig. 3a) using population genomic analyses of double-digest Restriction site Associated Sequences (ddRADseq; Barker *et al.* 2017). Here we use single nucleotide polymorphism (SNP) information from these ddRADseq reads to quantify genome-wide sequence divergence among genotypes from these four populations (*N*= 155 individuals; Supporting Information Table S1; Dryad#). Detailed methods for sequencing and SNP generation are described in Barker *et al.* (2017). Briefly, total genomic DNA was isolated using a CTAB/PVP DNA extraction protocol (Webb & Knapp 1990), and digested with enzymes *PstI* and *MseI* to create fragments for ddRADseq (Peterson *et al.* 2012). Unique combinations of individual P1 and P2 barcoded adapters were annealed to each sample, and resulting libraries size selected for fragments 350−650 bp. Size-selected libraries were enriched using 14 PCR cycles and sequenced on an Illumina HiSeq 2000 or 2500 to generate 100 bp paired-end reads. Reads were quality-filtered and de-multiplexed using SNOWHITE 2.0.2 (Dlugosch *et al.* 2013). R1 reads were trimmed to a uniform length of 76 bp for final SNP analysis. The denovo_map.pl pipeline program in STACKS 1.20 (Catchen *et al.* 2011, 2013) was used to merge identical reads into ‘stacks’, identify polymorphic sites, create a catalog of loci across individuals, and determine the allelic state at each locus in each individual (Barker *et al.* 2017). The ‘populations’ module in STACKS was used to export polymorphisms for analysis, and a locus was required to be genotyped in at least 70% of the samples in each population (-r 0.7), and have a minimum stack depth (-m) of ten. This resulted in 1585 polymorphic ddRAD loci (Dryad#). To quantify genetic variation, we calculated intrapopulation nucleotide diversity (π) as the average number of nucleotide differences among alleles (including invariant sites) for each ddRADseq locus using the ‘populations’ module in STACKS. To quantify divergence between two populations, we calculated pairwise interpopulation π (i.e. Dxy) between alleles at the same locus from each population. A custom script was used to combine polymorphism data from all loci to obtain π across the total length of all sequence tags.

### Experimental crosses

Experimental crosses were conducted to compare the performance of matings within and among the four native populations, using parents reared in a common environment (Supporting Information Fig. S1). We reared parents to flowering in a glasshouse at the University of Arizona (as in Dlugosch *et al.* 2015b). Between 9 and 22 parents / site were reared from seeds of different field mothers, or from bulk collections at Asian sites (N = 332; Supporting Information Table S1). Flowering heads (capitula) were covered with fine mesh bags while in bud, and hand pollinated using a single pollen donor when a large fraction of florets were receptive. Strong self-incompatibility in this species was verified by manual self-pollination and by bagging unmanipulated capitula (yielding 0% seed set). Seeds were collected at maturity from a total of 349 successful crosses within and among populations (Dryad#).

Progeny (N = 523, including 1−3 per cross) were reared for growth measurements in the common glasshouse environment. Increased growth is a fundamental metric of heterosis in experimental crosses (Birchler *et al.* 2010; Chen 2013), and it is a key trait whose evolution is associated with increased fitness in invasions of *C. solstitialis* (Eriksen *et al.* 2012; Dlugosch *et al.* 2015b). All size measurements of both ‘source’ genotypes (produced by within-population crosses) and ‘admixed’ genotypes (produced by among population crosses) were made in the same experiment, at both 4 and 5 weeks of age (Supporting Information Fig. S1). Size at both timepoints was measured using a non-destructive size index [(maximum leaf length * maximum leaf width)^1/2^ * leaf number] that has been shown to have a strong linear correlation with total biomass under glasshouse conditions in this species (Dlugosch *et al.* 2015b). Exponential growth rates between these two measurements for each plant were compared using REML Analyses of Variance (ANOVA) with fixed effects of 1) the source population of crossed genotypes; 2) observer (the person measuring the plants); and nested effects of 3) cross direction (the source population of the maternal vs. paternal genotype) and 4) individual parental combination. Least squares means (LSM) and standard errors were extracted from these models for use in analyses below.

### Mid-parent trait values

Source populations can be genetically divergent from one another in growth rate due to local adaptation or genetic drift, so we tested for evidence of non-additive genetic interactions during admixture by comparing admixed genotypes to mid-parent expectations that assume additivity in the trait (Dlugosch *et al.* 2015b). To calculate these expectations, both parental and admixed genotypes must be products of crosses conducted in a common environment, to minimize transgenerational plasticity effects on the traits of interest. Moreover, it is essential to compare measurements of parental and admixed genotypes at the same life stage within the same experiment, because growth rate is highly sensitive to experimental conditions. To accomplish this for *C. solstitialis*, we generated a distribution of pseudo-mid-parent values by randomly drawing combinations of ‘source’ growth phenotypes from the progeny of within-population crosses (i.e. representatives of parental lineages, produced by crosses in the common glasshouse environment, as described above) that were reared and measured at the same time as admixed genotypes (Supporting Information Fig. S1). Pseudo-parental combinations were drawn with replacement 1000 times, and their average (mid-parent) growth rates calculated to create a distribution of pseudo-mid-parent values that were compared to growth rates of admixed genotypes using two-tailed t-tests.

### Relationship between progeny performance and parental population diversity and divergence

To examine the relationship between growth rates in our experimental crosses and both genetic variation within source populations and genetic divergence between source populations, we used a linear model to explain deviation of growth patterns from mid-parent expectations (*Ymp*). The model included fixed effects in the form:

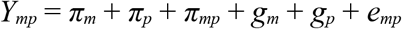

where π_*m*_ and π_*p*_ are maternal and paternal source intrapopulation π respectively, which are predicted to scale negatively with growth deviation due to the benefits of admixture at low within-population variation; π_*mp*_ is interpopulation π between parental source populations, predicted to influence growth deviation positively where genetic load is increasingly rescued, and negatively where there are epistatic incompatibilities accumulating at high divergence; *g_m_* and *g_p_* are respectively maternal and paternal source population LSM growth rate phenotype (‘source’ lineages grown in the same experiment as admixed progeny, as described above), predicted to correlate with growth deviation positively where there are transgenerational plasticity/epigenetic effects; and *e_mp_* is the residual error. All interaction terms were non-significant (*P* > 0.1) and are not shown. All non-significant terms were removed from the final model.

### Results

#### Literature Review

In total, we acquired data from 157 studies, which included introductions of 167 species (107 animals, 50 plants, 7 fungi, and 3 protists; Dryad#). Admixture was observed in 65 (39%) of these species, including only two cases where an introduction to one region had evidence of admixture whereas an introduction of the same species to another region did not (see Dryad#). The primary basis for determining admixture included formal tests among models of historical demography (11%), formal tests of population genetic signatures of admixture (12%), patterns of elevated genetic diversity (5%), similarity between invasions and potential source populations (85%), and population structure analyses (22%). Almost half of the studies (48%) utilized more than one of these methods as direct support for hypotheses for admixture, and only three studies inferred admixture solely from population structure analyses (see Dryad#).

Values of genetic differentiation among sample sites and diversity within these sites varied widely in the native range of both admixed and non-admixed introduced species (Fig. 2a). F_ST_ values spanned two orders of magnitude, with most values between 0.1 and 0.2 (Fig. 2a). Heterozygosity and haplotype diversity values spanned nearly the full range of 0 to 1, with median values in the region of 0.4 to 0.7 reflecting substantial variation within populations for most species (Fig. 2b). Species that demonstrated admixture in their introductions had lower median differentiation and higher median diversity than species without admixture observed, though these differences were not significant for any metric other than H_E_ (and see below that these overall differences may be influenced by marker type).

**Fig 2.**
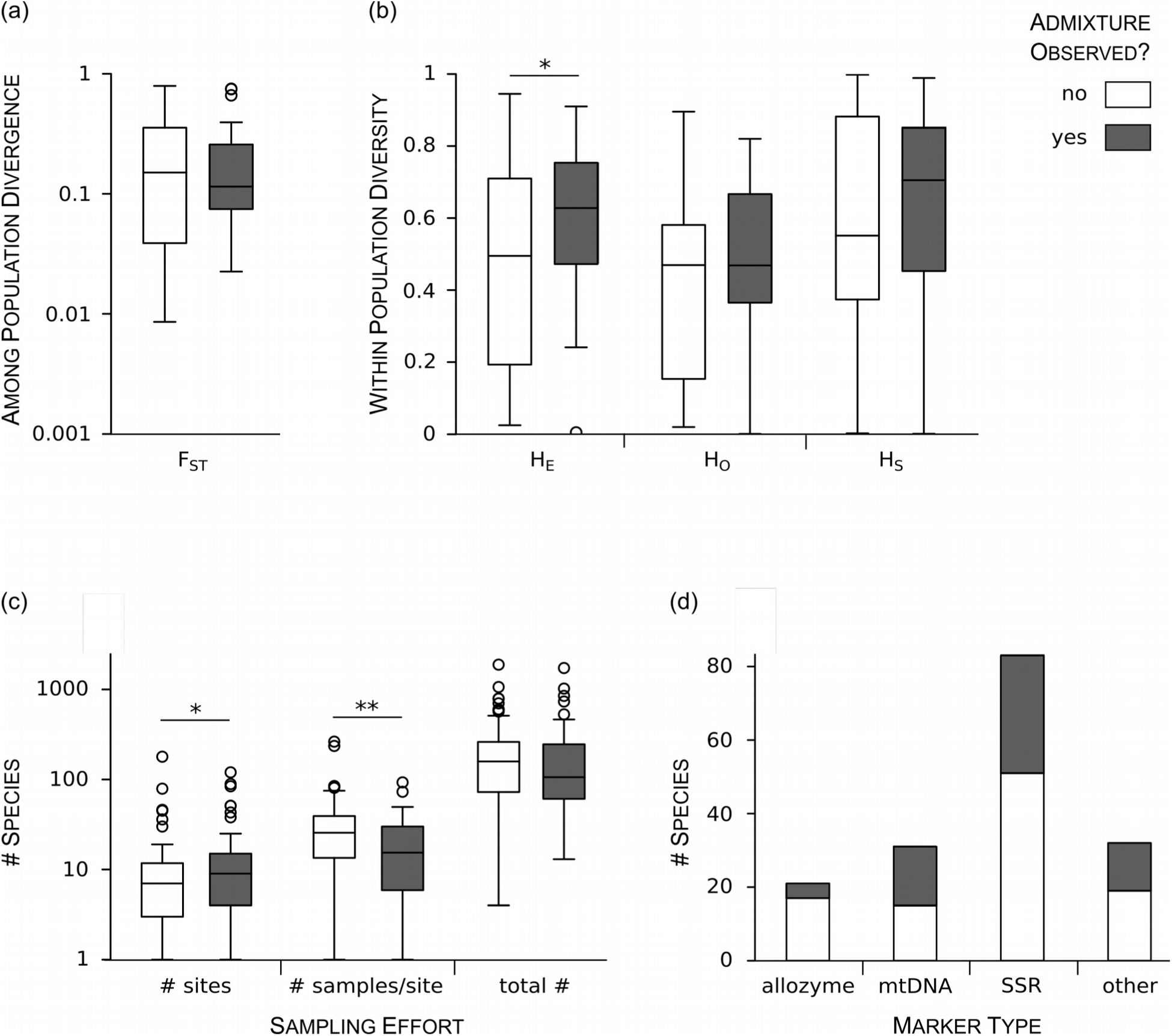
Descriptive statistics associated with observations of admixture (dark boxes) and no admixture (white boxes) in introduced species. (a) Divergence (FST and related metrics) among populations in the native range for each species, for studies using nuclear markers. (b) Average genetic diversity within populations in the native range for each species, as measured by HE and HO for nuclear markers and HS for cytoplasmic markers. (c) Sampling effort as the number of native sites, average number of samples within a site, and total native range sample number for each species. (d) The number of species surveyed with allozyme, mitochondrial DNA (mtDNA), microsatellite (SSR) and other (AFLP, ISSR, RAPD, ITS, nDNA) markers. \**P* > 0.05; \*\**P* > 0.01 for Wilcoxon Rank-Sum tests.

Several aspects of study design appeared to influence reports of admixture in introduced populations. Studies reporting admixture sampled 29% more sites in the native range and 40% fewer individuals within sites than studies not reporting admixture (Fig. 2c). The total number of individuals sampled across the native range did not differ significantly between studies that reported admixture in introductions and those that did not (Fig. 2c; Wilcoxon Rank-Sum test *P* = 0.14). In logistic regressions, the likelihood of reporting admixture was positively related tothe numbers of sites sampled in the native range *P* = 0.04 on ln transformed values) and negatively related to within site sampling effort (*P* = 0.001 on ln transformed values).

Marker type was also important for detecting admixture (Fig. 2d). The most commonly encountered genetic markers in our dataset included microsatellites (83 species), mitochondrial DNA (31 species), and allozymes (21 species). As might be expected, studies based on allozymes reported admixture less frequently (19%) than those based on more variable microsatellite and mitochondrial DNA, which reported admixture at least twice as often (38.6% and 51.6%, respectively). There was a marginal statistical effect of these common markers types on the likelihood of detecting admixture (contingency test: N = 135, *P* = 0.05). Differences among markers were not explained by sampling bias: the number of sites sampled in the native range did not differ among studies using these three different marker types (Kruskal-Wallis rank test: *P* = 0.70).

We examined the relationship between reports of admixture and metrics of genetic divergence and within-population genetic variation across native sites in microsatellite-based studies, which comprised just over half of our dataset. We examined these relationships using logistic regressions with the number of sites sampled as a covariate (ln transformed). We found no monotonic relationships between admixture and either genetic divergence (ln transformed values: model *P* = 0.93), or within-population diversity measured as H_E_ (model *P* = 0.42; Fig. 1d) or H_O_ (model *P* = 0.82; Supporting Information Fig. S2). Inspection of the data revealed a strong curvilinear relationship between admixture and genetic divergence (Fig. 1c), which did not appear to be explained by the number of sites sampled (Supporting Information Fig. S3).

#### Case Study: Experimental Crosses in C. solstitialis

Nucleotide diversity (π) varied within and across native *C. solstitialis* populations, and was consistently higher between pairs of populations than within populations (Fig. 3). Intrapopulation π ranged from 0.004 to 0.005 average SNPs/site, with the highest value in western Europe and the lowest in Asia. Interpopulation π increased to 0.005−0.008 average SNPs/site. The largest values of interpopulation π occurred in comparisons of alleles from southern Greece to those from other populations, consistent with our previous observation of ahighly differentiated lineage occupying the Apennine-Balkan Peninsulas (Barker *et al*. 2017).

**Fig 3.**
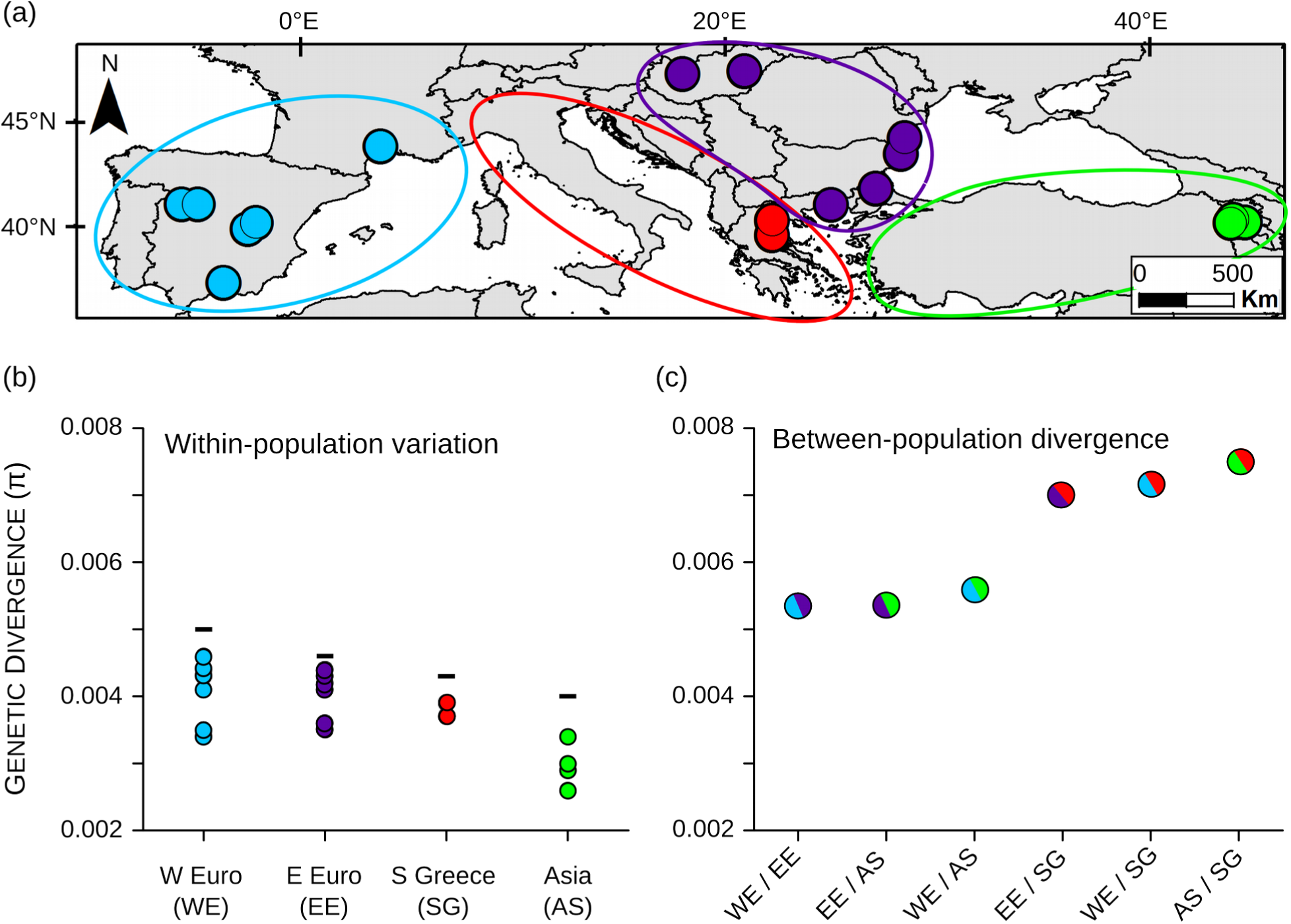
Sampling sites and sequence variation in the native range of *C. solstitialis*. (a) Sampling sites for this study (large dots) span genetically divergent populations in western Europe (WE, blue), eastern Europe (EE, purple), southern Greece (SG, red), and Asia (AS, green) as previously identified from population genomic analyses (previous sampling indicated by large and small dots; Barker *et al.* 2017). (b) Average intrapopulation nucleotide diversity (π) across the total length of all ddRADseq reads within each sampling site (dots) and for all individuals pooled within a region (bars). (c) Average interpopulation π in pairwise comparisons between individuals from different populations.

Growth rates differed significantly among different admixture combinations of parental source populations (F_5,11_ = 3.59, *P* = 0.005), spanning an order of magnitude in exponential growth rates (Supporting Information Fig. S4). Of the 12 combinations of crosses between genotypes from different maternal and paternal source populations, we found that seven deviated significantly from pseudo-mid-parent expectations, and all but one of these were in the positive (heterotic) direction (Fig. 4). Progeny of the maternal source population from western Europe experienced multiple heterotic interactions with those from other populations. The single negative interaction occurred in crosses with a maternal genotype originating from eastern Europe and a paternal genotype originating from the highly divergent population in southern Greece. In contrast, maternal genotypes originating from southern Greece showed positive genetic interactions with paternal genotypes from other regions, including those from eastern Europe.

**Fig 4.**
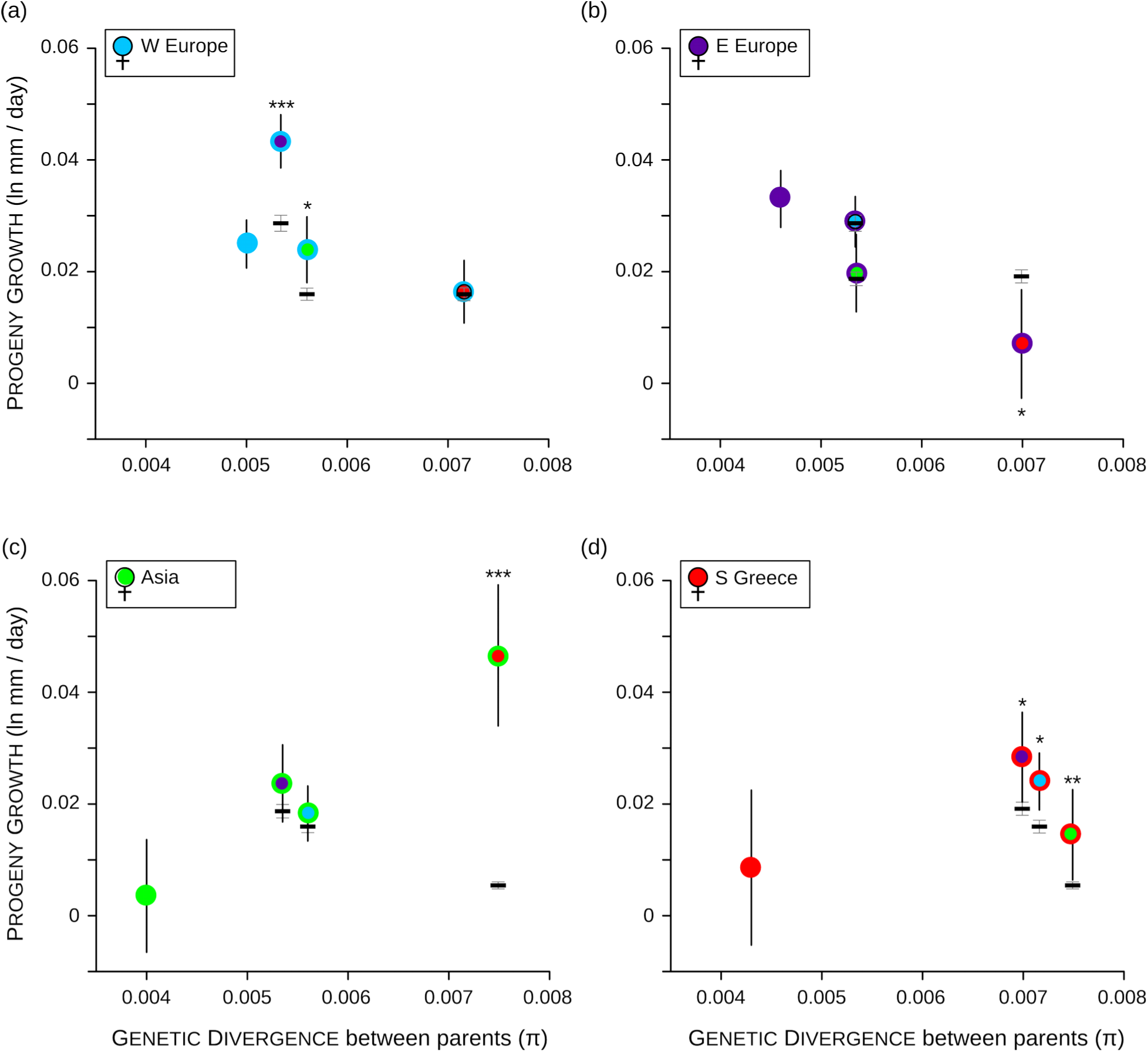
For each maternal region, panels (a−d) show growth rates of *C. solstitialis* progeny from experimental crosses to fathers from other regions and the mid-parent expectations for each cross. Growth rates are shown versus average pairwise nucleotide diversity (π) between parental populations. Color codes for each cross are as in Fig 2. Within-region crosses are indicated by solid colors, and between-regioncrosses by paternal region color outlined by maternal region color. Mid-parent expectations are shown as means (black bars) +/− s.e.m. (gray bars). Growth rates are least squares means +/− s.e.m., and significant deviations from mid-parent distributions are indicated as: *** *P* < 0.0001; ** *P* < 0.001; * *P* < 0.05.

Holding maternal source population constant, crosses to increasingly divergent paternal source populations showed a wide variety of trends in growth performance (Fig. 4), including positive (Asia), negative (eastern Europe), and curvilinear relationships (peaking at intermediate values; western Europe and southern Greece). Using data from all crosses, the deviations of admixed progeny from additive expectations were not predicted by parental source population divergence (interpopulation π), parental source intrapopulation π, or parental lineage phenotypes in a linear model (model *P* = 0.24). Removing the extreme heterotic datapoint in the cross of Asia × southern Greece (see Fig. 4a, Supporting Information Fig. S5), however, yielded a highly significant model, strongly predicting growth deviation (r^2^adj = 0.92; F_4,10_ = 28.6, *P* = 0.005), in which the direction of main effects were most consistent with epistatic interactions among loci (Fig. 1e,f). In particular, deviations in growth were negatively associated with divergence between parental source populations (Fig. 1e; *P* = 0.006), and positively associated with maternal source intrapopulation π, such that high genetic variation in the maternal source population made heterotic interactions stronger (Fig. 1f; *P* = 0.0007).

Progeny performance depended significantly on the source of the maternal vs. the paternal genotype in the cross (Fig. 5a; nested effect of cross direction: F_6,5_ = 0.014, *P* = 0.02). This result could suggest that transgenerational maternal effects influenced the growth of progeny, in which case a positive relationship would be predicted between maternal lineage phenotype and progeny deviation from mid-parent expectations. Yet, maternal lineage growth rate negatively predicted deviation from mid-parent expectations (Fig. 5b; *P* < 0.0001), and its interaction with genetic divergence was also negative (*P* = 0.0009). This result is not consistent with maternal effects, but could be explained by non-additive genetic interactions between the maternally-inherited cytoplasmic genome and the bi-parentally inherited nuclear genome.

**Fig 5.**
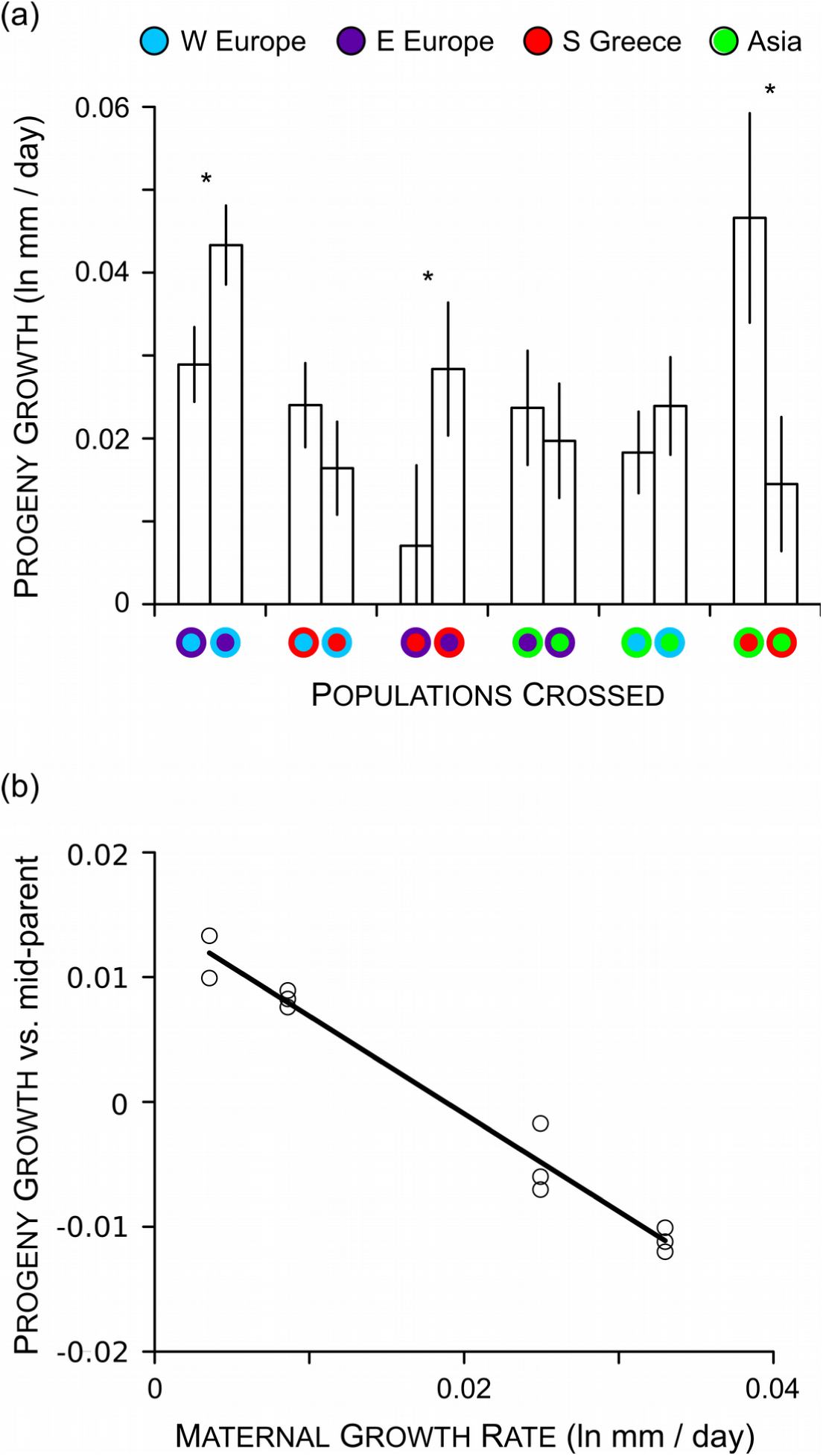
(a) Growth rates of *C. solstitialis* progeny from experimental crosses with all possible combinations of mother and father population-of-origin. Color codes for each cross indicate maternal and paternal population as in Fig 2, with paternal region color outlined by maternal region color. Growth rates are least squares means +/− s.e.m., and significant differences between reciprocal crosses of the same parental populations are indicated * *P* < 0.05. (b) In a linear model of progeny performance (excluding heterotic outlier [Asia × southern Greece]), growth rate of genotypes from the maternal region had a significant negative relationship (*P* < 0.0001) with the deviation of progeny from mid-parent expectations. Data in (b) show progeny deviation from mid-parent as partial residuals after taking into account all other effects in the model.

## Discussion

Abundant evidence for admixture in species introductions has raised the possibility that many invaders gain fitness advantages from associated increases in genetic diversity and novel genetic combinations. The benefits of admixture are expected to rise with increasing divergence between and decreasing genetic variation within source populations of introduced species, under a variety of mechanisms. We examined patterns of genetic variation reported in the literature and found a wide range of genetic differentiation and variation among potential source populations. Admixture was most prevalent at low to intermediate values of divergence among native populations, and was unrelated to levels of genetic variation within those populations. In our case study, admixture was generally beneficial in controlled crosses between *C. solstitialis* populations, but these benefits declined as divergence among populations increased and appeared to be affected by epistatic cytonuclear interactions. Our results suggest that admixture is frequent under conditions that are likely to be favorable for introduced species, but that these conditions may be bounded by negative genetic interactions at high levels of divergence.

### Literature Review

Our literature review supported the perspective that admixture is common in introduced species, appearing in almost 40% of the species studied. It is likely that this number is an underestimate; whether admixture was identified was sensitive to several aspects of power in the study design. First, studies that sampled a larger number of sites in the native range were more likely to identify admixture. This relationship is intuitive, given that many methods of detecting admixture rely on associating introductions with multiple potential sources, and this requires having sampled relevant source populations (Dlugosch & Parker 2008a; Cristescu 2015). Interestingly, sample size within sites was negatively related to observations of admixture. This could indicate a trade-off between sampling within site and the number of sites sampled, given that total sample size was relatively consistent. It is also possible that low sample size within sites inflates differences among potential sources, making admixture easier to detect, or creating the erroneous appearance that introductions are mixtures of what are inaccurate allele frequency estimates. Finally, inferences of admixture were sensitive to the resolution afforded by marker type, with admixture reported least frequently for allozymes,which tend to have low variation, and most frequently for microsatellites, which tend to have high variation). These effects of study design imply that admixture is likely to be difficult to detect where power is low, i.e. at low levels of divergence among source populations and at low within-population diversity (Dlugosch *et al.* 2015a). Indeed admixture was observed less often at the lowest levels of both of these factors in our review.

With regard to within-population diversity, introduced species in general showed a wide range of values in their native range, with relatively few species at the low end of the distribution where we might expect admixture to be most beneficial. Admixture tended to be observed in species with higher levels of heterozygosity overall, but this pattern could be driven by low power to detect admixture among populations with low marker diversity, and was not apparent within microsatellite-based studies (the most common type in our dataset). Several other factors may also obscure our ability to interpret whether introduced species are often in a situation to benefit from an increase in genetic diversity. Heterozygosity and haplotype diversity are less sensitive measures of genetic variation than metrics that better include rare alleles, such as allelic richness. On the other hand, heterozygosity should reflect the history of standing variation in a population (Lohr & Haag 2015), and rare alleles, by virtue of being rare, will not strongly influence most of the mechanisms that underlie the effects of admixture (with the notable exception of Evolutionary Rescue). We also emphasize that we have measured diversity in potential source populations rather than in founding populations that would have experienced admixture, but this information is unattainable for already admixed populations. Previous surveys have indicated that genetic bottlenecks are not typically severe during founding events, such that founder population genetic diversity largely resembles native population diversity, though this is not be true for all invaders and expansion fronts (Dlugosch & Parker 2008a; Excoffier *et al.* 2009; Uller & Leimu 2011; Dlugosch *et al.* 2015a). For example, several studies have found evidence consistent with inbreeding depression in natural invading populations and its rescue by admixture (e.g. Bailey & McCauley 2006; Nolte *et al.* 2009; Mullarkey *et al.* 2013; Keller *et al.* 2014; Rius & Darling 2014; van Kleunen *et al.* 2015). It may be that founding populations with particularly low genetic variation and high genetic load, which would be predicted to benefit most from admixture, are relatively rare across species introductions as a whole, although admixture should provide substantial benefits to founding populations when severe genetic bottlenecks do occur.

Potential source populations also demonstrated a wide range of genetic divergence estimates, but in this case admixture was disproportionately observed at low to intermediate values. This pattern could be explained by multiple processes. At low values of divergence, the positive relationship with increasing divergence could reflect increasingly favorable conditions for admixture to occur, due to the increasing fitness benefits of introgression predicted under all underlying mechanisms (Fig. 1a,b). As noted above, however, admixture is also likely to be underestimated at lower levels of divergence due to low power. The decline in observations of admixture at the highest levels of divergence, where the ability to detect sources of admixture should be strong, suggests that introgression between increasingly divergent sources is becoming less likely. This could reflect either 1) a lack of multiple introductions of divergent sources into the same location, 2) pre-zygotic isolating mechanisms that prevent introgression of additional divergent introductions, and/or 3) post-zygotic barriers to introgression (i.e. low fitness of admixed progeny), which select against introgression of additional divergent introductions. We consider each of these mechanisms in turn.

How multiple introduction sources tend to be distributed across a native range, particularly as a function of the genetic divergence of those sources, has not been investigated to our knowledge. Multiple introductions from different parts of the native range appear to be frequent across species introductions (Dlugosch & Parker2008a; Uller & Leimu 2011). Nevertheless, it seems probable that geographic barriers will constrain which source regions supply introduced propagules in some cases, and that these barriers might align with strong barriers to gene flow, such as large mountain ranges or bodies of water. The correspondence of introduction and genetic barriers could lead to a lack of admixture at high genetic divergence among native populations. This possibility could be studied given knowledge of specific source populations for both failed and successful multiple introductions, as well as the geographic distribution of genetic variation within the species involved. Such datasets might be available (or possible to accumulate) for groups with particularly well-documented introduction attempts, such as birds (e.g. Maitner *et al.* 2012).

Given multiple introductions of divergent material, prezygotic isolation among particularly divergent source populations could prevent the formation and establishment of admixed genotypes (Rius & Darling 2014). Though typically associated with isolation between species, high ecological divergence can also occur among conspecific populations (Hereford 2009). Genetic and ecological divergence among populations can reduce the likelihood of introgression through mechanisms such as differences in reproductive phenology, morphology, and behavior, or via inbreeding mating systems. For example, a recent study by Wadgymar and Weiss (2016) experimentally explored the introgression of a divergent plant population into an established population, and estimated a 40-90% reduction in admixed matings as a result of phenological mismatch.

Finally, admixture is expected to be unfavorable at high divergence among source populations due to low fitness of admixed progeny under epistasis. This pattern is a general prediction of epistatic interactions among loci that have been diverging among populations (Carroll *et al.* 2003; Phillips 2008), though epistasis is rarely mentioned in the context of invasions (Rius & Darling 2014). Negative epistatic interactions are expected to increase in their fitness effects as populations diverge, ultimately resulting in reproductive isolation and speciation (Orr & Turelli 2001). Epistasis is the most likely basis of the many observations of outbreeding depression (poor performance of among-population matings) within species, and non-native species would be exceptional if they did not also experience these interactions.

Other authors have pointed out that admixed progeny might also suffer from the loss of local adaptation, due to introgression of a divergent source that is not as well adapted to local conditions (Verhoeven *et al.* 2011). In order to explain the negative relationship that we found between reports of admixture and genetic divergence, the degree of ecological divergence between sources would have to be related to their degree of genetic divergence, which is often not the case (McKay & Latta 2002). On the other hand, negative epistasis would be predicted to result from divergence due to both local adaptation and genetic drift. In the absence of negative epistatic interactions, it becomes much more likely that natural selection will favor admixed progeny that retain locally adapted alleles, while benefitting from other positive effects of admixture, and introgression at some level will occur. Invaders might also be uniquely poised to avoid problems with loss of local adaptation from the native range, given that they occupy a novel environment (Rius & Darling 2014). Experimental studies of admixture across realistic levels of population divergence will be critical for disentangling these mechanisms, and revealing whether genetic interactions shape the success of matings among sources (Hufbauer 2017).

### Case Study: C. solstitialis

Significant genetic interactions were common in our crosses among native *C. solstitialis* populations, and all but one were positive (heterotic) in the first generation. Notably, some of the strongest evidence of heterotic interactions (with some of the highest resulting plant growth rates) occurred when the western European population served as the maternal parent. This lineage appears to be a primary contributor of invasions into the Americas (Eriksen *et al.* 2014; Barker *et al.* 2016). Strong heterotic interactions in this lineage suggest that a possible origin of successful invasion ‘bridgeheads’, which initiate many subsequent invasions (Lombaert *et al.* 2011), could be their propensity to foster beneficial admixture events (e.g. Turgeon *et al.* 2011).

The only significant negative interaction occurred when eastern European maternal genotypes were crossed with paternal genotypes originating from southern Greece. The population in southern Greece is particularly divergent from other populations (Fig. 2c; Barker *et al.* 2017), and may belong to a distinct subspecies. The geographic boundary separating southern Greece and eastern European populations may be a region in which early speciation dynamics, including the evolution of reinforcement, might fruitfully be studied. These particular interactions raise intriguing questions about which lineages are serving as the maternal parents in admixture
events between differentiated populations in the native range and in the invasions of *C. solstitialis*.

Both the identity of the maternal parent population of origin and genetic diversity within the maternal population were important to the performance of *C. solstitialis* crosses, which strongly suggests that the divergence of maternal (cytoplasmic) DNA from the paternal nuclear genome is the underlying cause of the interactions that we observed. There is increasing evidence that cyto-nuclear interactions within species are common and can have important phenotypic effects in both animals and plants (Ballard & Melvin 2010; Bock *et al.* 2014). The additional effect of genetic diversity with the maternal population would seem to suggest that maternal heterozygosity in some way reflected the potential for positive interactions between maternally inherited cytoplasmic genes and nuclear genes from a divergent paternal population. We are not aware of an established mechanism that could account for this pattern, though it seems plausible that a recent evolutionary history with a greater diversity of nuclear backgrounds might pre-dispose the maternal cytotype to have favorable interactions with novel paternal genotypes. Intriguingly, crosses between maternal genotypes from the invaded range in California (USA) and paternal genotypes from the invasion’s origin in Spain showed reduced seed set in a previous study (Montesinos *et al.* 2012). Compared to other populations across the species range, those from western Europe and California have some of the lowest levels of genetic divergence between them, but the highest phenotypic divergence (Barker *et al.* 2017), suggesting that adaptation might be driving the accumulation of negative epistatic interactions in this case.

### Epistasis and invasion biology

Perhaps because epistasis has rarely been discussed as a mechanism of major importance to admixture’s role in invasions, there have been few previous tests for epistatic interactions in introduced species. A small number of empirical studies have found evidence of epistasis underlying both negative and positive interactions. Keller and colleagues (2000) crossed native populations of three widespread agricultural weeds and found evidence of negative epistatic interactions in either the first generation (F1) cross or the backcross (F2) in each. Johansen-Morris and Latta (2006) crossed two invading genotypes of *Avena barbata* in their introduced range and found evidence for epistasis underlying both hybrid vigor in early generation hybrids (F2) and reduced fitness in later (F6) generations. Notably, later generations were highly variable and some individual lines showed potential for outperforming parental genotypes, revealing rare opportunities for beneficial admixture even when there is an overall pattern of deleterious hybrid breakdown.

Epistasis is unique in its potential to generate both positive and negative interactions during admixture. Any positive interactions are expected to diminish and become negative in later generations, as recombination breaks up co-adapted allele combinations. Thus benefits from epistasis during admixture are expected to be transient over time. Experiments with first generation crosses should generally demonstrate the maximum benefit of epistasis, and will be conservative with respect to identifying negative interactions that might ultimately constrain interbreeding of divergent populations. Importantly, transient increases in fitness can be highly beneficial when founding populations are struggling to establish (i.e. the ‘catapult effect’; Drake 2006). Further, negative epistatic interactions from hybrid breakdown in later generations can also be avoided by backcrossing with resident parental populations (Dlugosch *et al.* 2015b), and high fitness of early generation admixed genotypes can be maintained through clonal propagation or polyploidy in some cases (e.g. Facon *et al.* 2008). Indeed, clonal or polyploid spread of hybrids have provided some of our best examples to date of the evolution of invasiveness (Ellstrand & Schierenbeck 2000; hufbauer 2008). The complexity of these outcomes from epistasis alone argues that we should expect admixture to vary in its contribution to the establishment and invasion of introduced populations.

### Conclusions

Early discussions of admixture in species introductions recognized its potential to provide genetic and evolutionary rescue, and favorable genetic interactions (Novak & Mack 1993; Ellstrand & Schierenbeck 2000). These benefits were hailed as the resolution to what was sometimes called the ‘genetic paradox’ of invasions, wherein invaders were somehow successful despite experiencing founding events that were expected to have unfavorable effects on genetic diversity and fitness (Allendorf & Lundquist 2003; Kolbe *et al.* 2004; Hufbauer 2008). Particularly given that multiple introductions seemed to be quite common in successful invasions, admixture offered the potential to resolve the genetic paradox for many if not most invaders. We now know that introduced species don’t often suffer from large reductions in genetic diversity during founding events (Dlugosch & Parker 2008a; Uller & Leimu 2011; Dlugosch *et al.* 2015a). Nevertheless, our literature review does support the idea that admixture is common in species introductions, and that many species should have opportunities to benefit from its positive effects on fitness.

Importantly, we found that admixture is disproportionately lacking from species with high divergence among native populations, suggesting that the conditions for beneficial admixture are constrained. We note that negative epistatic interactions among genotypes should be expected at high divergence and could explain this pattern. Our case study of *C. solstitialis* demonstrated heterotic interactions in many crosses among native source populations, but also decreasing fitness benefits and ultimately negative interactions at high divergence, likely due to the action of cyto-nuclear epistasis. Additional experimental investigations across many taxa will be essential to understanding the mechanisms shaping the fitness effects of admixture in natural systems. Also needed are studies of systems with known introduction attempts, wherein it is possible to identify opportunities for admixture, whether those opportunities have been realized, and whether the prevalence of admixture is associated with the degree of fitness benefits. In general, our analyses argue that conflicting evidence for the benefits of admixture during invasion could be examined in the context of genetic divergence between and diversity within source populations to establish whether these different outcomes might be generally predictable, and therefore whether we should realistically expect that admixture is a general mechanism for the success of introduced species.

## Acknowledgements

We thank MS Barker and A Guggisberg for seed collections; K Gibson, C Patterson, and SW Smith for assistance with plant propagation and measurements; LK Honeker and S Tran for assistance with genotyping; A Zeeger for greenhouse logistics; and # reviewers for helpful comments on drafts of this manuscript. This study was supported by the National Institute of General Medical Sciences of the National Institutes of Health under Award #K12GM000708through the Center for Insect Science at UA to BSB, USDA ELI Fellowship #2017-67011-26034 to JEB, and USDA grant#2015-67013-23000 to KMD. The content is solely the responsibility of the authors and does not necessarily represent the official views of the National Institutes of Health.

## Data Accessibility

- Data from the literature review are available at Dryad#
- Data from experimental crosses are available at Dryad#

## Author Contributions

KMD conceived the study. SRA, JEB, FAC, HDG, and KMD performed the literature review. BSB performed the genomic analyses. JEC performed the experimental crosses. BSB and KMD analyzed the data and wrote the manuscript, which was edited by all authors.

## Supporting Information

Supporting information includes a single PDF document.

